# An Efficient and Safe Trans-complementation System for MPXV Mimicking Authentic Viral Infection

**DOI:** 10.1101/2023.12.29.573498

**Authors:** Jianying Liu, Longchao Zhu, Lingling Mei, Yuanyuan Liu, Yuanyuan Qu, Yulin Yuan, Fenfen Zhang, Yingyi Cao, Yibin Zhu, Wanbo Tai, Jun Ma, Min Zheng, Xiaolu Shi, Yang Liu, Gong Cheng

**Author notes:** These authors contribute equally (Jianying Liu, Longchao Zhu).

## Abstract

Since the Mpox virus (MPXV) outbreak in 2022, there have been 97,745 cases and 203 fatalities. This outbreak features increased transmission efficiency and a higher infection rate in the MSM population, with the underlying causes remaining unknown. The requirement of BSL-3 laboratories poses a barrier to MPXV research and antiviral development. Here, we report an efficient and safe trans-complementary system that produces a single-round infectious MPXV, preserving the virus’s authentic architecture and enabling it to complete its life cycle in complementing cells. This deficient MPXV genome, lacking two essential genes crucial for late gene transcription and viral assembly, is restricted to a single-round infection in non-complementing cell lines. Notably, SCID mice inoculated with this deficient MPXV exhibited no detectable disease or viral load. This deficient MPXV platform has demonstrated its capacity to study innate immunity and cell death during infection in complementing cells. It can also be used for antibody neutralization assays and anti-MPXV drug evaluation. This trans-complementation platform, safe for use in low-biosafety laboratories, offers a valuable resource for MPXV research and countermeasure development.

## Introduction

In the first half of 2022, the Mpox virus (formerly monkeypox virus, MPXV) suddenly emerged in multiple countries and rapidly spread. To date, a total of 97,745 laboratory confirmed cases and 660 probable cases, including 203 deaths, have been reported to WHO from 116 countries^1^. The primary natural hosts of the MPXV are mammals such as rodents, and it mainly spreads through close contact. Symptoms of MPXV range from fever, headache, and rash to severe complications like pneumonia and sepsis, potentially leading to death^2^. The MPXV has two primary genetic evolutionary branches: the West African clade with a 1% fatality rate and the Congo Basin clade with a higher fatality rate of up to 10%^3^. Evidence indicates the 2022 MPXV outbreak is genetically linked to the 2017-2018 Nigerian Mpox epidemic, suggesting it originated from the ongoing evolution of that strain^4^. The MPXV genome, composed of double-stranded DNA, typically evolves at a slow pace. However, the 2022 epidemic strain has surpassed the expected mutation rate, indicating an accelerated evolution of the virus^5^. These adapted mutations may alter the infectivity and transmission capabilities of the current epidemic MPXV. The widespread transmission of MPXV among MSM (Men who have Sex with Men) populations also indicate a potential change in tissue and organ preferences^6,7^. However, due to the lack of efficient and safe reverse genetic tools for MPXV, there is no definitive biological evidence to support these claims. The functional and phenotypic changes associated with these mutations, as well as the underlying causes and molecular mechanisms, have not been clearly elucidated.

The MPXV is enveloped, approximately 200-300 nm in diameter, brick-shaped, and features a dumbbell-shaped core^8^. The genome of the MPXV is characterized by a large, double-stranded DNA structure, approximately 197 kilobases in length. The MPXV genome encodes for numerous proteins, involved in various aspects of the virus’s life cycle, replication, and interaction with the host’s immune system. During the life cycle of the MPXV, two forms of viral particles are present: Intracellular Mature Virus (IMV) and Extracellular Enveloped Virus (EEV). The IMV possesses a single layer of the cellular membrane, whereas the EEV gradually acquires a double-layered phosphatidylethanolamine membrane through a trans-Golgi transport process. Both forms of viral particles are infectious^9^. The Mpox virus infection within cells progresses through early, intermediate, and late stages. Upon entering the cells, both IMV and EEV release prepackaged viral proteins and enzymatic factors into the cytoplasm, stimulating expression of early genes. Early protein synthesis triggers DNA replication, and production of intermediate transcription factors, followed by expression of late genes, mainly structural proteins. Eventually, virions assemble from DNA concatemers, incorporating all the necessary components for initiating a new infection cycle^10^.

Until now, several infectious clone systems of Orthopoxvirus genus have been established, including vaccinia virus (VACV)^11^, horsepox virus (HPXV)^12^, cowpox virus (CPXV)^13^. The trans-complementation deficient virus system represents a novel viral platform for the studies of high pathogenic viruses under low biosafety conditions^14^. This deficient virus lacks several essential genes necessary for viral replication and propagation, allowing for progeny production of the deficient virus only within specific cell lines that complement these essential genes^14^. To generate the trans-complementation system of MPXV, we selected the G9R and A1L genes on the MPXV, based on the previous research of Orthopoxvirus genus viruses. The G9R gene encodes a late transcription factor VLTF-1, while A1L gene encodes the late transcription factor VLTF-2, both are highly conserved across poxviruses^15^. These two genes have been proven essential for infection and replication within the poxvirus family; loss of either one of them results in the lethality of the poxvirus^16,17^. The locations of these two critical genes, widely separated on the genome, further reduces the risk of viral homologous recombination reverting to the wild type.

In our study, we chemically synthesized 56 Mpox viral genome fragments and assembled them into the complete viral genome *in vitro*, omitting two essential genes, G9R and A1L, and incorporating two fluorescence reporters under the intermediate (G9R) and late (A46R) gene promoters. Supported by fowlpox virus (FWPV), the modified MPXV genome with a hairpin structure was transfected into Vero E6 G9R+A1L complementing cells to rescue the live virus^11^. We assessed the pathogenicity and stability of this deficient MPXV, confirming its safety in various cell types and SCID mice^18^. Notably, the deficient MPXV was shown to elicit standard immune responses and cell death in complementing cell lines. Additionally, the integrated reporter genes proved useful for antibody neutralization tests and high-throughput drug screening. Our results indicate that this MPXV trans-complementation system is safe for low-biosafety conditions, providing a significant tool for MPXV basic and applied research.

## Results

### A trans-complementary deficient MPXV system

The trans-complementation system for MPXV produces deficient viral particles capable of only a single round of infection in non-complementing cells^14^. This system includes: (1) viral genomic DNA without the essential G9R and A1L late transcription factors, crucial for viral vitality, and (2) a complementing cell line that continuously expresses the G9R and A1L proteins. The 197 kb MPXV genomic DNA from the MA001 strain (ON563414.3) was segmented into 56 smaller fragments, chemically synthesized individually. These fragments were then assembled into seven larger fragments and inserted into the pSMART plasmid for amplification. Subsequently, these seven fragments were digested using Type II restriction enzymes and ligated to construct the complete MPXV genome *in vitro*. This deficient viral genome omits two essential genes, G9R and A1L, and features a mNeonGreen fluorescence gene at the beginning of the non-essential A46R gene, linked via a 2A linker, alongside a mCherry fluorescence gene under the G9R gene promoter (termed as MPXV G9R+A1L-KO) (Fig. 1a). Successful ligation of the seven large fragments was confirmed by PCR targeting the discontinuous regions between each of these fragments (Extended Data Fig. 1a). The hairpin structure, essential for Orthopoxviruses, was chemically synthesized and its structure integrity was verified with Mung bean nuclease treatment (Extended Data Fig. 1b). Hairpins were ligated to both ends of the viral genome to generate a functional genomic DNA construct, as confirmed by Sanger sequencing across the ligation site.

**Figure 1.**
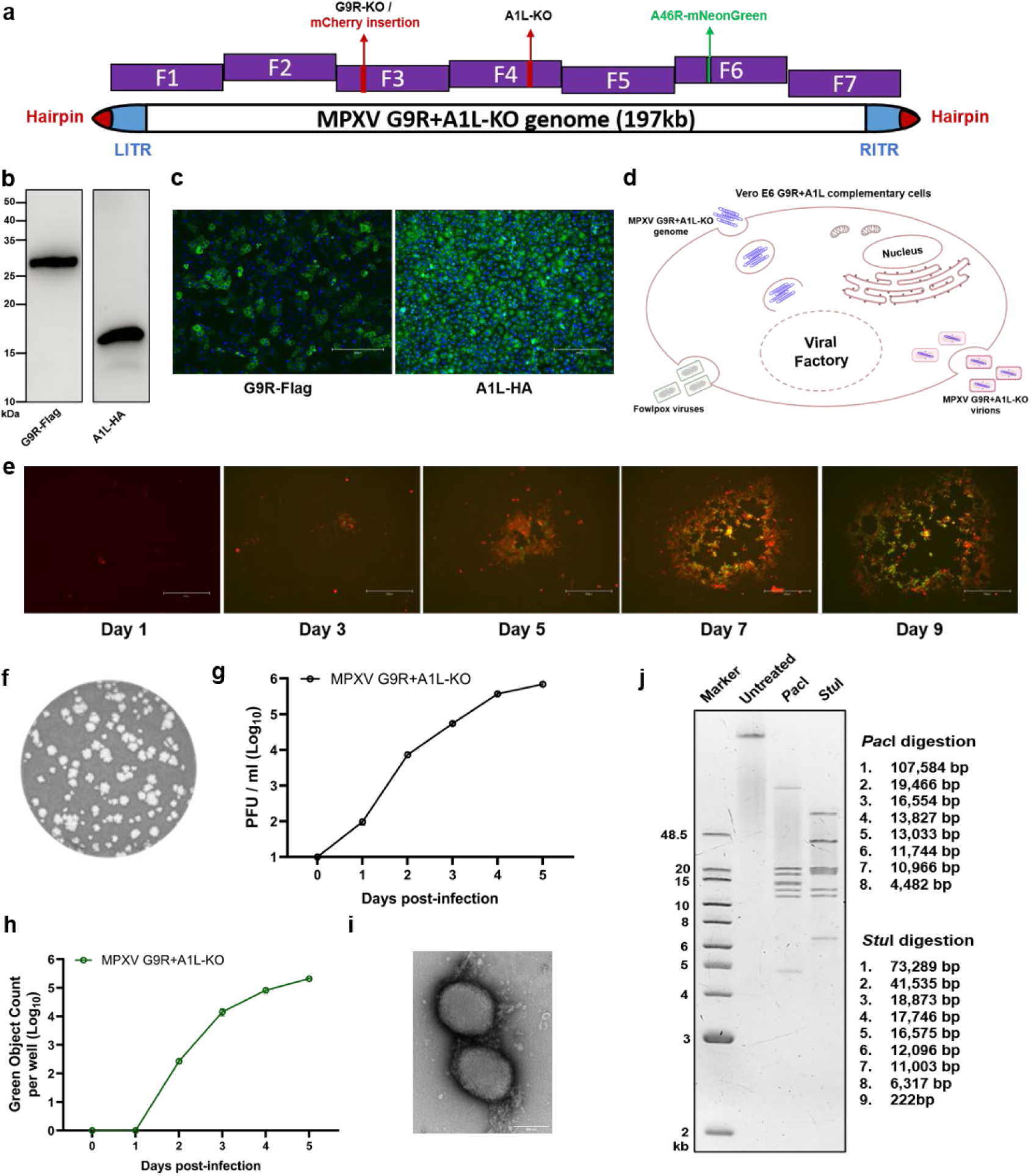
Construction and rescue of trans-complementation deficient Mpox virus. **a,** Schematic representation of deficient MPXV G9R+A1L-KO construction. Deletions of the G9R and A1L genes, along with the insertion of fluorescent proteins, were introduced into the backbone of MPXV MA001 strain. The complete MPXV genome was assembled from seven big fragments. Red arrows highlight the loci of G9R and A1L gene knockouts on the MPXV genome, with an mCherry reporter inserted at the G9R site. The green arrow marks the insertion site of the mNeonGreen reporter. **b,c,** The expression and localization of G9R and A1L genes in Vero E6 G9R+A1L complementing cells were detected by western blot **(b)**, and immunofluorescence assay **(c)**; Scale bar, 300 µm. **d,** The diagram for rescue of deficient MPXV infectious clone. The Vero E6 G9R+A1L stable cell lines were pre-infected with FWPV two hours prior to transfection with the deficient MPXV G9R+A1L-KO genome. FWPV facilitated the initiation of early-stage RNA transcription in the deficient MPXV. Due to species-specific constraints, the FWPV failed to produce infectious viral particles in mammalian cells. Therefore, only the progeny of the deficient MPXV was reproduced after several days of incubation. **e,** Observation of time-lapse MPXV G9R+A1L-KO viral plaque formation using mCherry and mNeonGreen fluorescence detection; Scale bar, 300 µm. **f,** Plaque morphology of MPXV G9R+A1L-KO virus in Vero E6 G9R+A1L complementing cells. The experiment was conducted using a plaque assay. Briefly, after a 2-hour incubation with MPXV G9R+A1L-KO virus, the cells were washed with DPBS to remove unattached particles. An overlay medium containing carboxymethyl cellulose was added, and the cells were incubated for 6 days. Plaques were stained with crystal violet. **g,h,** Replication kinetics of MPXV G9R+A1L-KO virus in Vero E6 G9R+A1L complementing cells. 0.01 MOI of MPXV G9R+A1L-KO virus was inoculated onto Vero E6 G9R+A1L cells. Following a 2-hour incubation, the cells were washed three times with DPBS and continuously cultured for 5 days with fresh 2% FBS DMEM. The virus titers and mNG-positive cells were determined using a plaque assay (**g**) and fluorescence determination (**h**). **i,** Electron microscopy of MPXV G9R+A1L-KO virus particles with negative staining; Scale bar, 200 nm. **j,** The full-length genomic DNA or restriction enzyme-digested fragments of deficient MPXV were analyzed by pulse-field gel electrophoresis (PFGE). The expected sizes of the digested fragments were presented but fragments smaller than 2 kb will not be detected in this assay.

To produce complementing cells, the G9R and A1L genes (linked by IRES,internal ribosomal entry site) were incorporated into a lentivirus delivery system, which included puromycin or blasticidin for antibiotic selection. The Vero E6 (African green monkey kidney) cells were infected with recombinant lentivirus and after a two-week antibiotic selection, isolated Vero E6 G9R+A1L colonies were analyzed by western blot and immunofluorescence assay (IFA) to verify the expression of G9R and A1L genes (Fig. 1b,c). The Vero E6 G9R+A1L stable cell lines were then pre-infected with FWPV two hours prior to transfection with the deficient MPXV G9R+A1L-KO genome. FWPV facilitated the initiation of early-stage RNA transcription in the deficient MPXV, a process shared among all Orthopoxviruses. Due to species-specific constraints, the FWPV failed to produce infectious viral particles in mammalian cells. Therefore, only the progeny of the deficient MPXV was reproduced after several days of incubation (Fig. 1d). Typically, significant cytopathic effects (CPE) are noticeable 5 to 7 days after transfection (Fig. 1e). The cells are then harvested 10 days post-transfection, at which point significant cell death is observed in about half of the cell population. The deficient MPXV G9R+A1L-KO forms plaques in the Vero E6 G9R+A1L cells normally (Fig. 1f)^19^ and can effectively amplify in the complementing cells (Fig. 1g and Extended Data Fig. 2a), simultaneously generating a green fluorescence signal (Fig. 1h). Then, the deficient viral particles were purified through sucrose density gradient centrifugation and analyzed using western blot with an anti-MPXV A35R polyclonal antibody (Extended Data Fig. 2b). The negative staining of MPXV G9R+A1L-KO particles by a transmission electron microscope confirmed their structural integrity (Fig. 1i). To further verify the integrity of the viral genome, genomic DNA of MPXV G9R+A1L-KO was isolated from the aliquoted viral stock and digested with selected restriction enzymes. The resulting DNA fragments were then characterized by pulsed-field gel electrophoresis (Fig. 1j). These results demonstrate that the deficient MPXV G9R+A1L-KO was successfully rescued and efficiently reproduced in complementing cells, with a normal poxvirus structure and an intact full-length genome. Sequencing results indicate that the final rescued Mpox virus has two synonymous mutations in the coding region (A75900T and T182846A) and an 18 bp deletion in the non-coding region (179205-179222); otherwise, the sequence is completely accurate.

### The deficient MPXV exhibited sing-round infection in non-complementing cells

The MPXV G9R+A1L-KO virus, lacking two essential genes for viral reproduction, was restricted to single-round infections in non-complementing cells. It regains its reproductive capacity and produces progeny viruses only when infecting cells that fully complemented both G9R and A1L genes (Fig. 2a). To determine the lethality linked to these two genes, the MPXV G9R+A1L-KO was infected in cells partially complementing either the G9R or A1L gene (Fig. 2b). Notably, MPXV-G9R+A1L-KO failed to replicate in cells complementing only one gene and did not produce any green fluorescence four days post-infection (Fig. 2c,d). Subsequently, 14 non-complementing cell lines from various mammals, including animals and humans, and each derived from different tissues, were selected to assess the replication ability of MPXV G9R+A1L-KO. These cell lines were inoculated with a high initial multiplicity of infection (MOI) of 1 (Fig. 2e, f). Supernatants with cell lysates were collected daily for plaque assays. For the first two days, a small residual of the inoculated virus was detected, but no amplification occurred in any non-complementing cell line from day 3 to day 5 (Fig. 2e). The fluorescence detection clearly showed the absence of green fluorescence in non-complementing cells from day 1 to day 5, while the deficient virus replicated efficiently in the Vero E6 G9R+A1L complementing cells (Fig. 2f, Extended Data Fig. 3).

**Figure 2.**
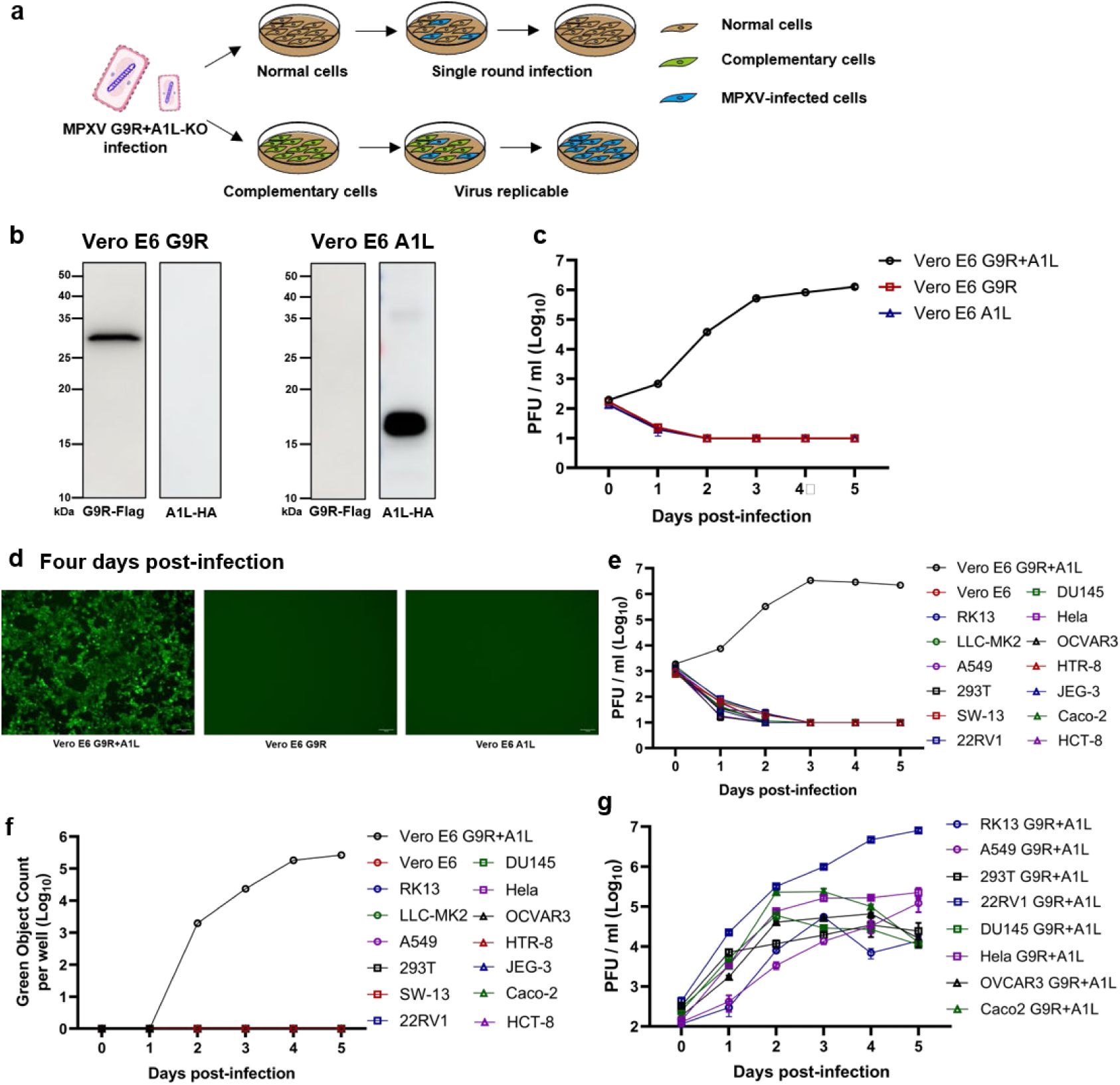
Replication dynamics of MPXV G9R+A1L-KO virion in non-complementing and complementing cells. **a,** Schematic representation of the replication of MPXV G9R+A1L-KO virion in both non-complementing and complementing cells. **b,** The expression of G9R and A1L genes in Vero E6 G9R and Vero E6 A1L partially complementing cells respectively. **c,** Replication kinetics of the MPXV G9R+A1L-KO virus in Vero E6 G9R+A1L, Vero E6 G9R, and Vero E6 A1L complementing cells. Each cell type was inoculated with 0.1 MOI of the MPXV G9R+A1L-KO virus. The virus titers were determined using a plaque assay. **d,** Fluorescence microscopy analysis of deficient MPXV infected complementing cell lines. Representative mNeonGreen (mNG) positive images at 4 days post infection are shown. Scale bar, 100 µm. **e,f,** Assessment of single-round infection of MPXV G9R+A1L-KO virus on non-complementing cell lines. 1 MOI of MPXV G9R+A1L-KO virus was inoculated onto 14 different cell lines. The Vero E6 G9R+A1L cell served as positive control. The virus titers and mNG-positive cells were determined using a plaque assay (**e**) and fluorescence determination (**f**). **g,** Replication kinetics of MPXV G9R+A1L-KO virus in different G9R+A1L complementing cells. 0.1 MOI of MPXV G9R+A1L-KO virus was inoculated onto eight G9R+A1L complementing cells. The virus titers were determined using a plaque assay. **c-g,** For different types of cells, the initial number of infected cells is normalized to an equal amount. **c,e,g,** The cells were lysed with the culture medium by freezing and thawing twice for plaque determination at the indicated times. The detection limit for the plaque assay is 10 PFU/ml.

### The deficient MPXV was reproducible in various complementing cell lines

Vero E6 cells are immune-deficient and lack the capability to produce interferon. Given this, we investigated whether MPXV G9R+A1L-KO could amplify in other G9R+A1L complementing immune-competent cell lines. Using the G9R+A1L lentiviral system, we introduced these two complementary genes into 8 representative cell lines. Following antibiotic selection, we assessed the expression of these two knocked-in genes by western blot. Blotting results confirmed the successful expression of the G9R and A1L genes in 8 cell lines, although expression levels varied among the different cells (Extended Data Fig. 4). Upon inoculating these stable cell lines with MPXV G9R+A1L KO, the virus rapidly amplified in all of them. The amplification showed significant variation, with the highest viral load observed in 22RV1 cells, derived from human prostate carcinoma epithelial cells (Fig. 2g). This result indicated that the MPXV MA001 strain may have a tissue preference for prostate carcinoma epithelial, which may correlate with the increased infection rates in MSM populations during this outbreak^6^. It should be noted that no correlation was observed between the viral load and the expression levels of complementary genes in the cells (Fig. 2g and Extended Data Fig. 4).

### The deficient MPXV remains stable throughout serial passages

To validate the genetic stability of these deficient viruses, we subcultured MPXV G9R+A1L-KO up to 20 passages (P20) on Vero E6 G9R+A1L cells in three independent replicates. Viruses from P5, P10, P15, and P20 were subjected to PCR analysis to verify the continuous absence of the targeted G9R and A1L genes and the stability of the mNeonGreen fluorescence gene (Extended Data Fig. 5a). All gene deletions and insertions remained stable after 20 passages (Extended Data Fig. 5b). Although the morphology of the plaques appeared slightly larger after 20 passages (Extended Data Fig. 5c), there were no significant green fluorescence changes four days post-infection between P1 and P20 viruses (Extended Data Fig. 5d). Only one mutation (Δ146999T) affecting protein characteristics were consistently detected across three replicates through full genome sequencing (Extended Data Table 1). Additionally, the genomic DNA of deficient MPXV could not be detected by qPCR in non-complementing cells after two blind passages (P2).

### Safety evaluation of deficient MPXV virions *in vivo*

To further evaluate the safety of deficient MPXV *in vivo*, we selected the SCID mice, which were susceptible to Orthopoxvirus infection. We inoculated these mice intraperitoneally with a maximum dose of purified MPXV G9R+A1L-KO virions, equivalent to approximately 10^6^ PFU/mouse. An initial titer of 2×10^5^ PFU/mouse of vaccinia virus (VACV, tiantan strain) was served as the positive control. Clinical symptoms and body weight were monitored, and blood were collected daily for viral load detection. All mice were sacrificed 9 days post-infection to analyze organ viral loads (Fig. 3a). The results indicated no weight change between the MPXV G9R+A1L-KO and Mock groups, compared to a notable weight loss in VACV-infected mice (Fig. 3b). Additionally, no viremia or clinical symptoms were observed in either the Mock or deficient MPXV groups (Fig. 3c,d). Nine days post-infection, no viral nucleotide residuals were detected in organs from the deficient MPXV group, while significant amounts of VACV were present in all organs (Fig. 3e). Pathological analysis revealed clear damage in the lungs and spleens of VACV-infected mice, but not in those from the deficient MPXV and Mock groups (Fig. 3f). These results demonstrate that deficient MPXV G9R+A1L-KO induces only a single-round infection and lacks virulence in mice.

**Figure 3.**
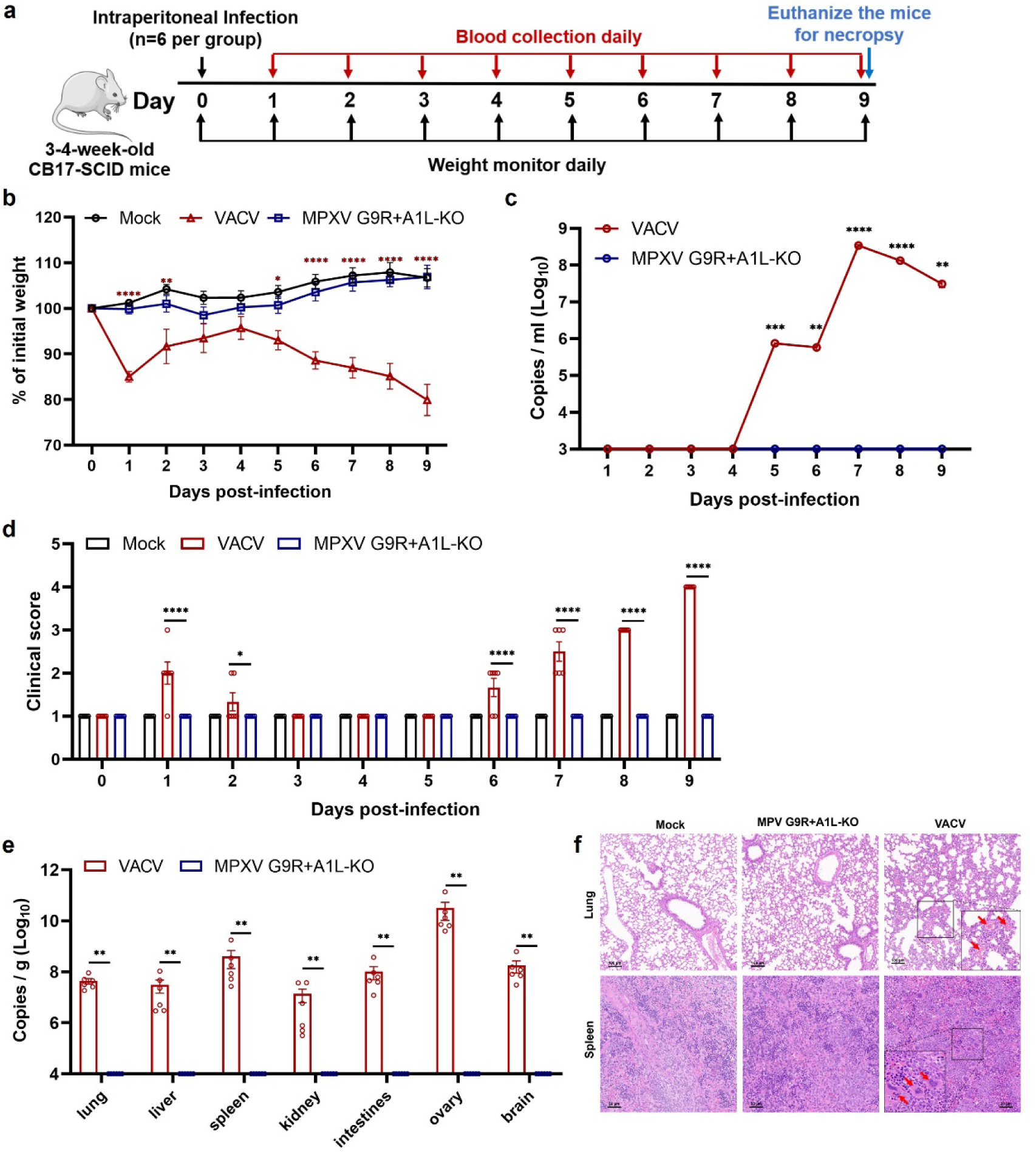
The safety evaluation of MPXV G9R+A1L-KO virion in animal models. **a,** Experimental scheme of MPXV G9R+A1L-KO virus challenge. 3-4 weeks old SCID mice were intraperitoneally (I.P.) inoculated with 1×10^6^ PFU of MPXV G9R+A1L-KO virus, 2×10^5^ PFU of VACV, or PBS mock control, respectively. Mice were monitored for weight loss, disease, and viral load. **b,** The weight changes of animals after intraperitoneal infection of MPXV G9R+A1L-KO virus (n=6) and VACV (n=6). The uninfected mock group (n=6) served as a negative control. Body weights were recorded daily for 9 days. Data are presented as mean ± standard deviation. Weight changes between MPXV G9R+A1L-KO and mock or VACV groups were assessed using two-way ANOVA with Tukey’s post hoc test. The red asterisk indicates a statistical difference between the mock and VACV groups. No significant difference was observed between mock and MPXV G9R+A1L-KO groups. **c,** Viral replication kinetics in serum of infected animals. The amounts of genomic DNA were quantified by quantitative PCR. **d,** Disease of MPXV G9R+A1L-KO and VACV-infected animals. The diseases include ruffled fur, lethargic, hunched posture, and orbital tightening. The clinical scores are presented. P values were adjusted using the Bonferroni correction to account for multiple comparisons. **e,** The viral loads in lung, liver, spleen, kidney, intestines, ovary and brain after infection with G9R+A1L-KO virus and VACV. Dots represent individual animals (n=6). **f,** Pathological observations of lung and spleen sections in mice. Representative hematoxylin and eosin (H&E) staining images were presented. **c-e,** The mean ± standard error of mean is presented. A non-parametric two-tailed Mann-Whitney test was used to determine the differences between VACV and MPXV G9R+A1L-KO groups. **b-e,** Differences were considered significant if *p* < 0.05; *, p < 0.05; **, p < 0.01; ***, p < 0.001; ****, p < 0.0001.

### The deficient MPXV regulates the innate immune response

The deficient MPXV retains the ability to modulate immune responses and activate other cellular signaling pathways, including programmed cell death, in complementing cell lines. MPXV G9R+A1L-KO was introduced at an MOI of 0.1 into IFN-α stimulated Hela G9R+A1L cells, with the RNA transcriptome measured at 48 hours post-infection. Results indicated that while IFN-α significantly upregulated interferon-stimulated genes (ISGs), MPXV showed a robust inhibitory effect on ISG expression (Fig. 4a), demonstrating similar transcriptome profiles to those in VACV-infected cells^20^. Meanwhile, like typical Orthopoxvirus infections, inoculation of MPXV G9R+A1L-KO in Hela G9R+A1L cells induced slow but apparent cell death over time, with nearly 20% of cells dying at 48 hours post-infection (Fig. 4b,c). The activation of apoptosis was indicated by the cleavage of caspase 3, 7, and 8 in deficient MPXV-infected cells (Fig. 4d). These results suggest that this trans-complementation system is a valuable platform for studying innate immune and inflammatory responses, as well as cell death, during MPXV infection *in vitro*.

**Figure 4.**
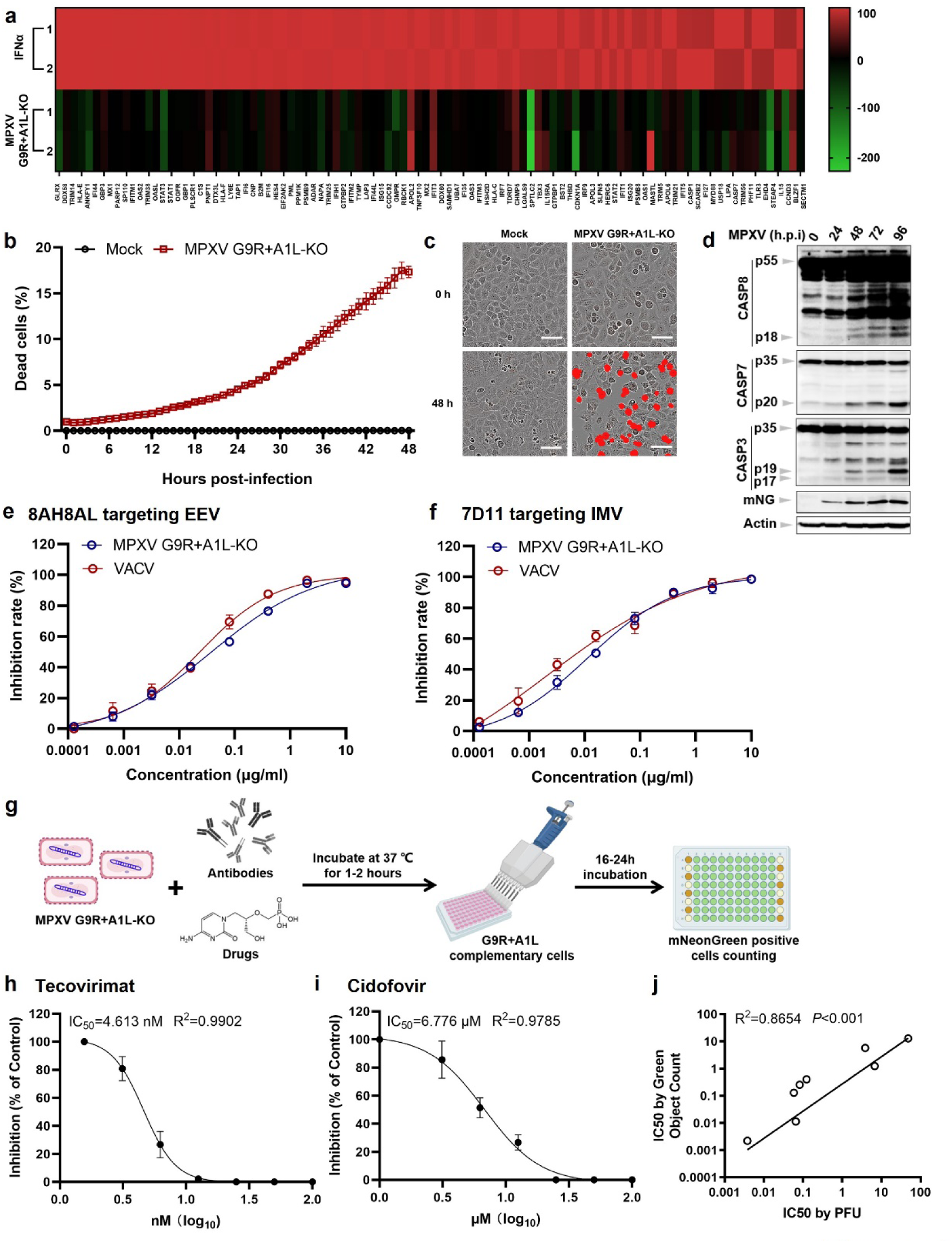
The potential of deficient MPXV for pathogenesis studies, antibody neutralization and high-throughput drug screening. **a,** Heatmap of differentially expressed genes (DEGs) associated with IFNα responses in Hela G9R+A1L cells. The heatmap shows the relative percentages of DEGs involved in IFNα responses in deficient MPXV-infected samples versus uninfected cells, with a color scale on the right indicating expression levels. **b-d,** The deficient MPXV stimulates the programmed cell death. **b,** Cell death kinetics were assessed using the IncuCyte live-cell automated system. Hela G9R+A1L cells, seeded in 24-well plates, were infected with MPXV G9R+A1L-KO virus at an MOI of 0.1. Dead cells, stained with propidium iodide (PI), were counted hourly over a 48-hour period. **c,** PI-positive cells were marked with a red mask for visualization. Scale bar, 80 µm. **d,** The activation of caspase 3, 7, and 8 post-MPXV G9R+A1L-KO infection at different time points. **e,f,** Neutralization activity of monoclonal antibodies (mAbs) against MPXV G9R+A1L-KO and VACV. The 8AH8AL(**e**) and 7D11(**f**) antibodies were incubated with EEV and IMV forms of MPXV G9R+A1L-KO and VACV, respectively, for neutralization via a plaque assay. Curves were fitted using nonlinear regression and mean ± standard deviations from two independent experiments were shown. The experiment was conducted twice with similar results. **g,** Scheme diagram for high-throughput MPXV antibody or drug screening using an mNG positive cell counting assay. **h,i,** The IC_50_ assessment of tecovirimat and cidofovir potency against deficient MPXV in complementing cells. Vero E6 G9R+A1L cells were infected with MPXV G9R+A1L-KO virus and treated with indicated concentrations of tecovirimat (**h**) or cidofovir (**i**) for 6 days. Plaque formation inhibition is expressed in %, normalized over control conditions. IC_50_ and R^2^ are indicated. Data are presented as mean ± standard deviations. Experiments were performed twice in triplicate. **j,** Correlation analysis of IC_50_ values between the mNG positive cell counting assay and plaque formation assay. The Pearson correlation coefficient (R²) and p-values (two-tailed) are indicated.

### The anti-MPXV antibodies and drugs evaluation by deficient MPXV

Due to the extensive capacity of the MPXV genome, the deficient MPXV system can accommodate multiple reporters, including fluorescence proteins and luciferase etc. This allows for the establishment of a high-throughput system for screening and evaluating anti-MPXV antibodies and drugs. Firstly, we compared the antibody neutralization capabilities of two previously reported mAbs, 8AH8Al^21^ and 7D11^22^, against MPXV G9R+A1L-KO and VACV using the PRNT_50_ assay, the gold standard for antibody neutralization tests. These two antibodies neutralized MPXV G9R+A1L-KO as effectively as VACV, demonstrating that viral particles produced by the trans-complementation system are suitable for assessing antibody efficacy (Fig. 4e,f). Subsequently, we evaluated the potential of the deficient MPXV for high-throughput drug screening (Fig. 4g). MPXV G9R+A1L-KO was pre-mixed with various drugs previously reported to have anti-MPXV capabilities^23–25^. The anti-MPXV IC_50_ values for these compounds were then determined using the classical plaque-forming unit (PFU) assay and a high-throughput green fluorescence cell counting method, respectively. Notably, two drugs, tecovirimat and cidofovir, displayed potent anti-MPXV efficacy (Fig. 4h,i). The IC_50_ values measured by our trans-complementation system were consistent with those reported in previous studies using authentic live virus^23,24^. Furthermore, we compared the correlation of IC_50_ values derived from the PFU and fluorescence methods, finding that the results from both methods were comparable and correlated well (Fig. 4j, Extended Data Table 2). These findings suggest that this MPXV trans-complementation system is effective for high-throughput, automated screening and evaluation of antibodies and drugs.

## Discussion

Our reverse genetic trans-complementation platform for MPXV, along with various complementing cells, offers a powerful tool for *in vitro* studies on MPXV genetic evolution, life cycle, as well as the induced innate immune responses and programmed cell death. The assembly of the full-length deficient MPXV can be achieved by T4 ligation of seven individual fragments *in vitro*, offering several advantages over traditional recombination and single plaque selection methods^11^. This *in vitro* approach provides an alternative method for obtaining genetically modified MPXV, avoiding the complex plasmid transfer processes across yeast and bacteria. Additionally, multiple mutations or genetic modifications can be simultaneously introduced into the MPXV genome, eliminating the need for multiple rounds of single virus plaque screening and identification. It may also reduce the frequency of unexpected mutations and recombination compared to the *in vivo* assembly system^12^.

The primary advantage of this deficient MPXV system is its enhanced safety. For high-pathogenicity viruses, direct genetic modification of the wild-type virus could potentially enhance pathogenicity, raising biosafety and bioethical concerns. The trans-complementation system introduces an additional safety lock for reverse genetic engineering. The conditional replication in complementing cells confines the modified pathogens to these specific cells, significantly reducing risks to operators and the environment. Our results indicate that immune regulation and programmed cell death function normally in complementing cells with deficient MPXV, maintaining the capacity for MPXV-host interaction studies. Moreover, the viral structure is identical to the authentic virus, ensuring the system’s efficacy for antibody and drug screening while maintaining safety (Fig. 4).

Fowlpox virus was employed to trigger the initiate transcription of the MPXV G9R+A1L-KO DNA genome due to the conservation of polymerases with the Orthopoxvirus genus. Once entry into the cells, FWPV releases its RNA polymerase from the virion core, which then recognizes and initiates transcription and replication of the deficient MPXV genome. However, due to the absence and variation of specific accessory proteins, FWPV fails to assemble in mammalian cells and does not produce progeny viruses. his strategy ensures that only pure deficient MPXV particles are assembled and released, eliminating the need for single viral plaque selection and reducing the time required for mutant virus rescue. In cells infected with the produced deficient MPXV virus, the residual fowlpox virus genome was no longer detectable by qPCR.

In our trans-complementation system, we integrated two fluorescent proteins into the deficient MPXV genome. The mCherry gene, replacing the absent G9R gene, is regulated by intermediate transcription factors. The mNeonGreen gene was fused to the beginning of the late-expressed A46R gene. As a result, even without complementation of G9R and A1L genes, a high initial MOI infection of deficient MPXV in non-complementing cells can trigger red fluorescence, while green fluorescence appears only in cells where successful complementation occurs. Therefore, the red fluorescence can serve as a tool to evaluate viral single-round entry and replication, providing a quick and effective assessment approach in primary cells lacking complementation.

The Mpox virus genome does not enter the cell nucleus, resulting in a very low probability of recombining with the host cell genome. However, this trans-complementation system, while reducing safety concerns, still presents some risks due to MPXV’s double-stranded DNA nature, which could potentially recombine with other spontaneously infected poxviruses. To further minimize recombination risks, the two excised essential genes are broadly separated on the MPXV genome, and the complementary genes are codon optimized with different flanking nucleotide sequences. This spacing reduces the chance of reverting to a fully functional wild-type MPXV through a single recombination event. It should be noted that, although recombination between deficient MPXV and complementing genes in cells is unlikely, there is still a possibility that deficient MPXV could undergo long-segment recombination with other poxviruses to produce “chimeric viruses”. To mitigate this risk, co-infection of deficient MPXV with other mammalian poxviruses should be avoided. On the other hand, we tested complementing systems lacking three or more essential genes; however, they showed reduced reproductive efficiency. The two-gene knockout system represents the optimal compromise between safety and efficiency. Crucially, our extensive *in vitro* and *in vivo* safety evaluations affirm the adequacy of this two-gene knockout strategy.

Utilizing this trans-complementation deficient MPXV system, we can investigate specific MPXV gene functions by knocking out or modifying them, including introducing single amino acid mutations to pinpoint those critical for the 2022 MPXV epidemic. Concurrently, we can study crucial host genes influencing MPXV infection and reproduction using a CRISPR-Cas9 single-gene knockout library in complementing cells. Additionally, we can fuse fluorescent proteins to various non-essential MPXV genes to label different regions of the viral particle, such as the extra-membrane, intra-membrane, and core, allowing for precise tracking of cell attachment, entry, and movement via single-particle microscopy. Furthermore, we can generate custom-engineered viral tools for anti-MPXV antibody and drug screening and evaluation. Overall, our study offers a comprehensive platform for studying MPXV infection, deciphering disease progression, and developing countermeasures under low-safety conditions.

## Methods

### Ethics statement

Mouse studies were performed in accordance with the guidance for the Care and Use of Laboratory Animals of the Shenzhen Bay Laboratory (SZBL). The protocol was approved by the Institutional Animal Care and Use Committee (IACUC) at SZBL. All the animal operations were performed under anesthesia by isoflurane to minimize animal suffering.

### Animals and cells

CB17-SCID female mice were purchased from Syagen Biotechnology company (Suzhou, China). Vero E6 (CL-0491), Caco-2 (CL-0050), 22RV1 (CL-0004), RK-13 (CL-0501), LLC-MK2 (CL-0141), SW-13 (CL-0451A), DU 145 (CL-0075), A549 (CL-0016), 293T (CL-0005), Hela (CL-0101), OVCAR-3 (CL-0178), JEG-3 (CL-0127), HCT-8 (CL-0098), HTR-8 (CL-0765) cells were purchased from Procell (Wuhan, China) and cultured in high-glucose Dulbecco’s modified Eagle’s medium (DMEM, C11965500BT, Gibco) supplemented with 2 mM L-glutamine, 100 U/mL Penicillin-Streptomycin (P/S, 15140122, Gibco), and 10% fetal bovine serum (FBS; HyClone Laboratories, UT). For G9R+A1L complementing cell lines, the following concentrations of puromycin (ant-pr-1, InvivoGen) or blasticidin (ant-bl-1, InvivoGen) were supplemented respectively: Vero E6 G9R+A1L (puromycin, 1:500), RK13 G9R+A1L (puromycin, 1:4000), 22RV1 G9R+A1L (puromycin, 1:2000), Caco-2 G9R+A1L (puromycin, 1:1000), OVCAR-3 G9R+A1L (puromycin, 1:2000), A549 G9R+A1L (blasticidin, 1:100), DU 145 G9R+A1L (puromycin, 1:5000), 293T G9R+A1L (blasticidin, 1:200), and Hela G9R+A1L (blasticidin, 1:2000). All other culture media and supplements were purchased from ThermoFisher Scientific. All cell lines were authenticated using STR methods and tested negative for mycoplasma.

### Construction of the MPXV deficient infectious clone

For the construction of the deficient MPXV G9R+A1L-KO infectious virus, 56 DNA fragments covering the full-length MPXV genome (MA001 strain) were chemically synthesized by Qingke company. The deletions of the G9R and A1L genes, along with the insertion of fluorescent proteins, were introduced into the respective regions of genome (Fig. 1a). To generate the F1 to F7 sub-genomic fragments, seven to eight small DNA fragments were assembled the pSMART-BAC 2.0 (42030, Lucigen) plasmid backbone using Gibson Assembly (E2621L, NEB). These recombinant plasmids were then introduced into BAC-Optimized Replicator v2.0 electrocompetent cells (60210, Lucigen) via electroporation at 2500V, 25μF, 300Ω using the Bio-Rad GenePluser Xcell Total system (1652660, Bio-Rad). To construct the full-length MPXV G9R+A1L KO genome, the F1 to F7 plasmids were digested with the PaqCI enzyme (R0745L, NEB) and assembled *in vitro* using T4 ligase (M0202M, NEB) along with hairpins at both ends. The hairpins were chemically synthesized by Qingke company. The hairpin ssDNA sequence is 5’-CACATTTTTTTCTAGACACTAAATAAAATAGTAAAATATAATATTAATGTACTAAAACT TATGTATTATTAATTTATCTAACTAAAGTTAGTAAATTATATACATAATTTTATAATTAATTTAATTTTACTATTTTATTTAGTGTCTAGAAAAAAA-3’. The successful ligation of fragments was verified by PCR across the ligation sites and DNA sequencing. The ligation products were then introduced into Vero E6 G9R+A1L complementing cells to rescue the virus. The primers used for MPXV construction was listed in Extended Data Table 3.

### Selection of G9R+A1L complementing cell lines

To construct cell lines expressing the MPXV G9R+A1L proteins, the Flag-G9R and HA-A1L genes were chemically synthesized and optimized for mammalian codon expression. For packaging the lentivirus, the pLVX-G9R+A1L-puromycin or pLVX-G9R+A1L-blasticidin plasmids were transfected into 293T cells using the Lenti-X Packaging Single Shots kit (631275, TaKaRa). Lentiviral supernatants were harvested 48 hours post-transfection and filtered through a 0.45 μm membrane (SLHV033RB, Millipore, Burlington, MA). One day before transduction, target cells were seeded in a 6-well plate with DMEM containing 10% FBS. After 12-18 hours, cells were transduced with 0.5 mL of lentivirus for 24 hours in the presence of 12 μg/mL of polybrene (107689, Sigma-Aldrich, St. Louis, MO). Twenty-four hours after transduction, cells from a single well were distributed into four 10-cm dishes and cultured in medium supplemented with the designated concentration of puromycin or blasticidin. The medium containing antibiotics was refreshed every 2 days. After 2-3 weeks of selection, visible puromycin or blasticidin-resistant cell colonies were formed. Select colonies were moved into 24-well plates. Upon reaching confluence, cells were treated with trypsin and replated in 6-well plates for further growth. These cells were designated as P0 cells. For cell line validation, total cellular mRNA was extracted, subjected to RT-PCR, followed by cDNA sequencing of the G9R and A1L genes, and protein expression was verified via western blot.

### Rescue of the deficient MPXV G9R+A1L-KO infectious virus

The deficient MPXV G9R+A1L-KO infectious virus was recovered in the Vero E6 G9R+A1L cell line, which constitutively expresses G9R and A1L proteins. In brief, confluent monolayers of the Vero E6 G9R+A1L cell line in 12-well plates were infected with 1 MOI of avian pox virus (Quail adapted strain, CVCC AV1003) for 2 hours prior to MPXV DNA transfection. Subsequently, the culture media were replaced with Opti-MEM (31985062, Gibco), and DNA transfection was performed using Fugene 6 reagent (E2691, Promega). 1 μg of MPXV DNA was mixed with 3 μl of Fugene 6 reagent 100 μl Opti-MEM medium for 15 minutes at room temperature. The DNA mix was then slowly dripped onto the cells and gently mixed. Post 24-hour incubation, the cell supernatant was replaced with fresh 2% FBS DMEM for continued cultivation. Cell monitoring continued for 8-10 days until the appearance of cytopathic effects (CPE) and expression of mCherry and mNeonGreen fluorescent proteins. The supernatant and cells underwent 2 freezing and thawing cycles at −80 °C. After centrifugation at 1000 g for 10 minutes, the supernatants were aliquoted and stored at −80 °C as the P0 virus stock.

### Virus plaque assay

10-fold dilutions of the deficient MPXV virus stock were prepared in a 96-well plate in triplicate. The diluted MPXV viruses were added to the Vero E6 G9R+A1L cells for a 2-hour incubation. Cells were then washed thrice with DPBS to eliminate any unattached virus particles. 3 mL of overlay medium (2% FBS DMEM with 2% HEPES, 1% penicillin/streptomycin, and 0.8% carboxymethyl cellulose) was added over the cells for a 6-day incubation. Subsequently, cells were fixed with 4% formaldehyde for 1 hour and stained with 300 μL of 0.5% crystal violet solution for 5 minutes. Plaques were manually counted under whiteboard illumination.

### Virus replication kinetics determination

Virus replication kinetics were determined by using a plaque assay as described above. Typically, cell lines were seeded in 24-well plates and cultured at 37 °C with 5% CO_2_ for 16 hours. Deficient MPXV was then inoculated into the cells at the specified MOI and incubated at 37 °C for 2 hours. Post-infection, cells were washed thrice with DPBS to eliminate any unattached virus particles. At designated time points, supernatants and cells were collected and underwent 2 freezing and thawing cycles. Following centrifugation at 1000 g for 5 minutes, the supernatants were employed to determine virus titers via plaque assay.

### The extraction of MPXV genome

For virus DNA extraction, 5 mL of viral culture supernatant was treated with TURBO™ DNase (ThermoFisher, AM2238) for 30 minutes at 37°C, followed by RNase A (Sangon Biotech, B600473) for 30 minutes at 37°C. EDTA was then added, and the mixture was incubated at 65°C for 10 minutes to inactivate the DNase. The sample was mixed with virion lysis buffer (2 mg/mL Proteinase K, 6% sodium dodecyl sulfate, 250 mM Tris-HCl, pH 8.0, 20 mM EDTA, pH 8.0) and incubated at 37°C for 2 hours. The MPXV genome was gently extracted using the phenol-chloroform method and precipitated in three volumes of ethanol. Finally, the precipitate was collected by centrifugation at 15,000g at 4°C, rinsed with 70% ethanol, briefly dried, and resuspended in 20 μL of ddH_2_O.

### The pulse-field gel electrophoresis (PFGE)

For pulse-field gel electrophoresis (PFGE), full-length viral genomic DNA was digested with *Pac*I or *Stu*I for 2 hours at 37°C and separated using 1% Seakem Gold agarose gels (Lonza, 50150) prepared with half-strength Tris-borate-EDTA electrophoresis buffer. The DNA fragments were separated by the CHEF Mapper® XA system (BioRad) in Auto Algorithm mode (Molecular Weight: Low, set to 1 kb; Molecular Weight: High, set to 197 kb) for 9 hours at 14°C, using a switching time gradient of 1 to 10 seconds at 6.0 V/cm, a linear ramping factor, and a 120° angle. The DNAs were then visualized with SYBR Gold stain (ThermoFisher, S11494) and imaged using the ChemiDoc MP Imaging System (BioRad).

### Virus passage

The confluent monolayers of the Vero E6 G9R+A1L cell line in 6-well plates were infected with 0.1 MOI of deficient MPXV G9R+A1L-KO infectious virus with triple biological replicate. For 2-3 days, supernatants and cells were collected and underwent 2 freezing and thawing cycles. Following centrifugation at 1000 g for 5 minutes, the supernatants were aliquoted and stored at −80 °C as the P1A, P1B, P1C virus stock, respectively. 100μl of virus stock was transferred to 100% confluency of the virus producer cell line in the 6 well-plates for propagation. Whole genomic DNA sequencing was performed at every 5 passages by Nanopore sequencing. P0 and P20 sequences have been deposited to figshare website (Licence ID: CC BY 4.0, DOI: 10.6084/m9.figshare.26340355).

### Safety evaluation of deficient MPXV in mice model

Three-to-four-week-old female CB17-SCID mice were randomly assigned and intraperitoneally inoculated with 1×10^6^ PFU of MPXV-G9R-A1L-KO, 2×10^5^ PFU of VACV, or PBS mock control. Throughout the study, the mice were monitored daily for clinical signs. Blood samples were taken daily for viral DNA extraction to detect the presence of the virus. At 10 d.p.i., the mice were euthanized, and tissue samples were collected for hematoxylin and eosin (H&E) staining and virus titration. All samples were then stored at −80 °C until analysis.

### Real-time qPCR assay

Deficient MPXV was isolated using the EZNA SE Viral DNA/RNA kit (R6871, Omega) according to the manufacturer’s instructions. The absolute copy number of MPXV DNA was quantified using the 2×EasyTaq® PCR SuperMix (AS111-01, TransGene) by Taq-man qPCR. The detection probe and primers were based on the conserved DNA polymerase (F8L) gene and described as follows: Forward primer (5’-TCA ACT GAA AAG GCC ATC TAT G-3’), Reverse primer (5’-GAG TAT AGA GCA CTA TTT CTA AAT CCC A-3’), and probe 5’-FAM-CCA TGC AAT A(T–BHQ1) A CGT ACA AGA TAGTAG CCA AC-Phos-3’, both were chemically synthesized by Qingke company. The qPCR reactions were performed using the CFX96 real-time PCR detection system (1855195, Bio-Rad).

### Immunofluorescence assay and western blot

For immunofluorescence assay (IFA) staining, Vero E6 G9R+A1L cells were cultured on a Nunc™ Lab-Tek™ II Chamber Slide™ System. G9R and A1L proteins were labeled with primary anti-Flag (F1804, Sigma-Aldrich) and anti-HA (26183, Invitrogen), followed by secondary Alexa Fluor 488 anti-mouse HRP antibody (A28175, Invitrogen). The nuclei were then stained with DAPI. Fluorescence signals were captured using a Nikon fluorescence microscope. For western blot analysis, various types of cells transduced with G9R, A1L, G9R+A1L, or those infected with deficient MPXV were harvested and lysed using RIPA buffer. They were then probed with anti-Flag, anti-HA, and anti-MPXV A35R (RVV13101, AtaGenix) antibodies. The results were analyzed using ImageLab software version 6.0.1 (#12012931, Bio-Rad).

### Deficient MPXV antiviral drug evaluation

Vero E6 G9R+A1L cells were seeded in 12-well plates at a specified density. Twenty-four hours post-seeding, MPXV G9R+A1L-KO (with a final inoculation of 50 PFU per well) was pre-mixed with varying drug concentrations before adding to the cells. After a 2-hour incubation at 37°C and 5% CO_2_ with gentle rocking every 15 minutes, the inoculum was removed. Cells were then overlaid with a mixture of 2% HEPES, 1% penicillin/streptomycin, and 0.8% carboxymethyl cellulose (C104979, Aladdin) in DMEM, with each condition tested in triplicate. Plates were then incubated for 6 to 8 days at 37°C in a 5% CO_2_ incubator. Monolayers were fixed in 4% formaldehyde and stained with 0.5% crystal violet. Plaques were manually counted under whiteboard illumination. IC_50_ values were calculated using GraphPad Prism 9.

### Deficient MPXV neutralization assay

The neutralizing activity of monoclonal antibodies (mAbs) was determined using the EEV or IMV forms of VACV, or MPXV G9R+A1L-KO in a plaque reduction neutralization (PRNT) assay. Briefly, 100 PFU of either VACV or deficient MPXV were incubated with serial two-fold dilutions of the mAb for 30 minutes at 37°C and then added to monolayers of Vero E6 or Vero E6 G9R+A1L cells in 6-well plates. After a one-hour incubation, cells were overlaid with 0.8% carboxymethyl cellulose in DMEM with 2% FBS and incubated for 3 days (for VACV) or 6 days (for MPXV G9R+A1L-KO) at 37°C in 5% CO_2_ to allow plaque formation. Monolayers were then fixed with 4% formaldehyde and stained with 0.5% crystal violet. Plaques were manually counted under whiteboard illumination.

### Statistics

Mice were randomly allocated into different groups. The investigators were not blinded to allocation during the experiments or to the outcome assessment. No statistical methods were used to predetermine sample size. Descriptive statistics are provided in the figure legends. A linear regression model in the software Prism 9 (GraphPad) was used to calculate the IC_50_ values from the antiviral assay. Differences between continuous variables were assessed using the non-parametric Mann-Whitney test. The weight loss data are presented as mean ± standard deviation and were statistically analyzed using two-factor analysis of variance (ANOVA) with Tukey’s post hoc test. *, p < 0.05; **, p < 0.01; ***, p < 0.001; ****, p < 0.0001; ns, p > 0.05, not significant.

## Data availability

Extended Data includes 5 figures, and 3 tables can be found with this article. Any other information is available upon request. Mpox sequences have been deposited to figshare website (Licence ID: CC BY 4.0, DOI: 10.6084/m9.figshare.26340355).

## Acknowledgments

This study was supported by grants from the National Key Research and Development Plan of China (2023YFC2307400, 2021YFC2300200, 2021YFC2302405, 2022YFC2303200, 2022YFC2303400, and 2023YFC2305900), the National Natural Science Foundation of China (82241082, 32270182, 82372254, 32188101, 32273099, 82271872, 82341046 and 32100755), Shenzhen Medical Research Fund (B2301009), Shenzhen San-Ming Project for Prevention and Research on Vector-borne Diseases (SZSM202211023), Yunnan Provincial Science and Technology Project at Southwest United Graduate School (202302AO370010), the New Cornerstone Science Foundation through the New Cornerstone Investigator Program, and the Xplorer Prize from Tencent Foundation.

## Author contributions

G.C., Y.L. and J.L. conceived and designed the study. G.C., Y.L., J.L. and L.Z. wrote and revised the manuscript. J.L., L.Z., L.M., Y.L., Y.Q., Y.Y., F.Z., Y.C., Y.Z., W.T., J.M., M.Z., X.S. and Y.L. performed the experiments and analyzed the data. All the authors reviewed, critiqued, and provided comments on the text.

---

**Extended Data Figure 1.**
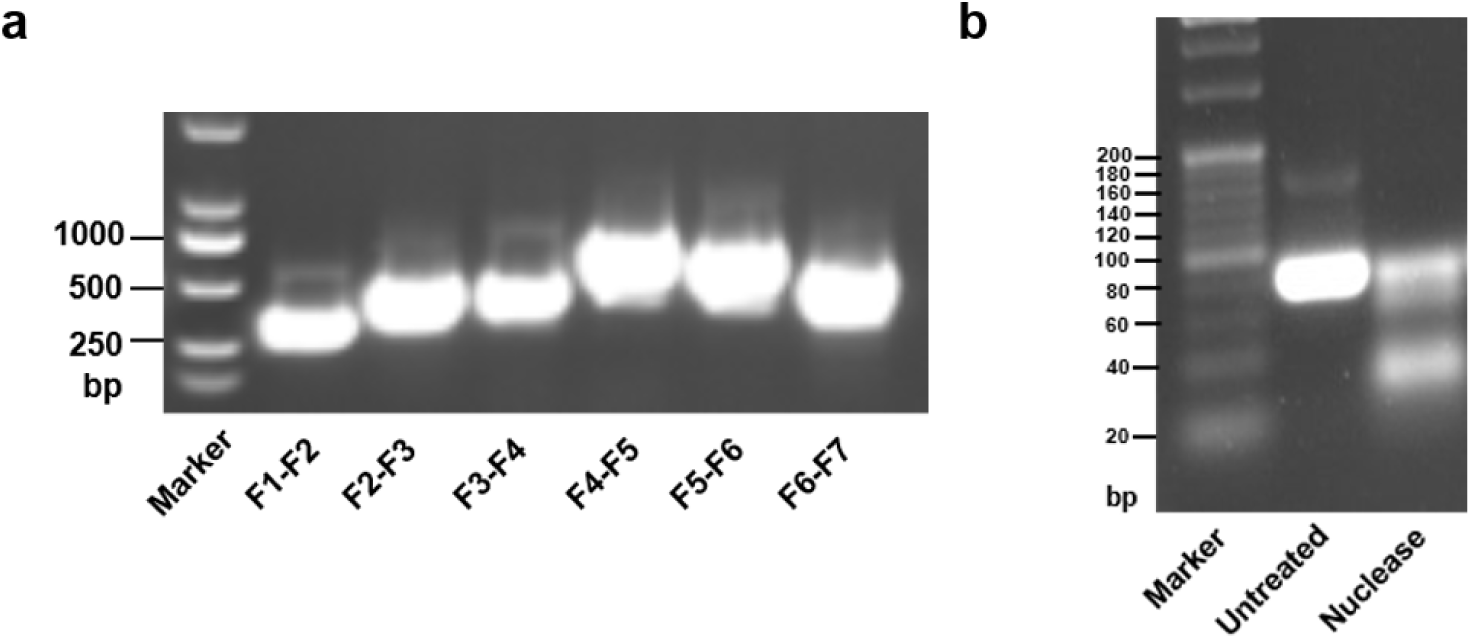
The validation of the integrity of the full-length MPXV genome and hairpin structures. **a,** Validation the integrity of full-length MPXV genome by PCR. Primers spanning fragment junctions were employed to verify the complete assembly of the MPXV genome. **b,** Validation of the secondary structure formation of chemically synthesized MPXV hairpin oligos. The Mung bean exo/endonuclease recognized and digested double strand DNAs.

**Extended Data Figure 2.**
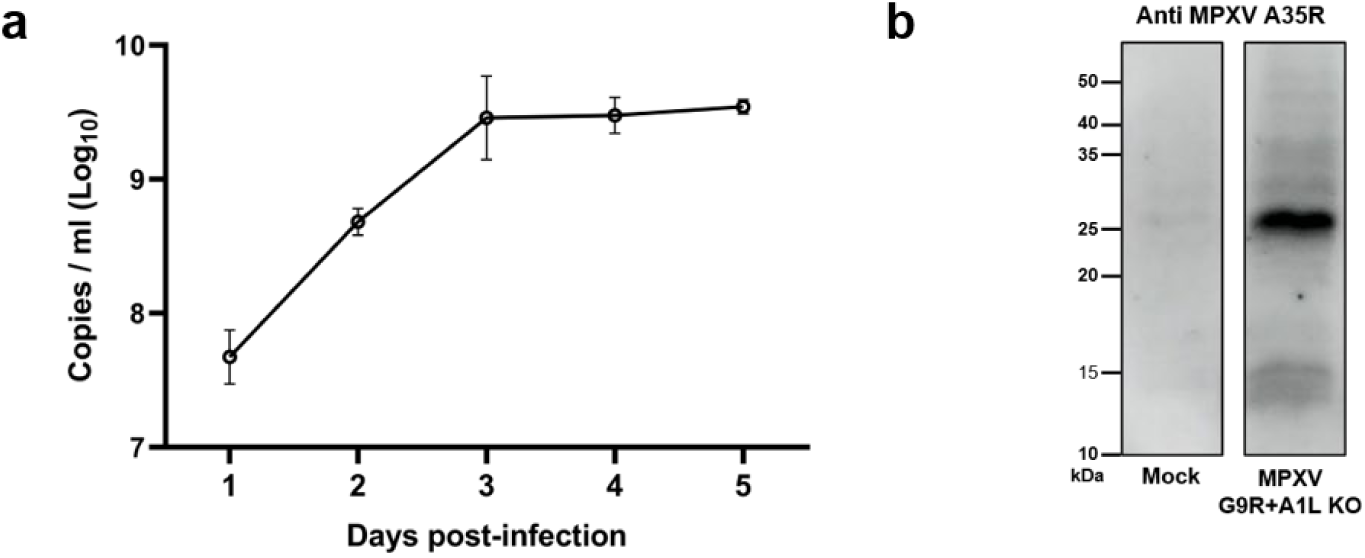
The qPCR and western blot verification of deficient MPXV. **a,** Viral replication kinetics of MPXV G9R+A1L-KO in Vero E6 G9R+A1L cells. The amounts of genomic DNA were quantified by quantitative real-time PCR. **b,** The expression of A35R gene in MPXV G9R+A1L-KO infected or un-infected cell lysates was detected by western blot.

**Extended Data Figure 3.**
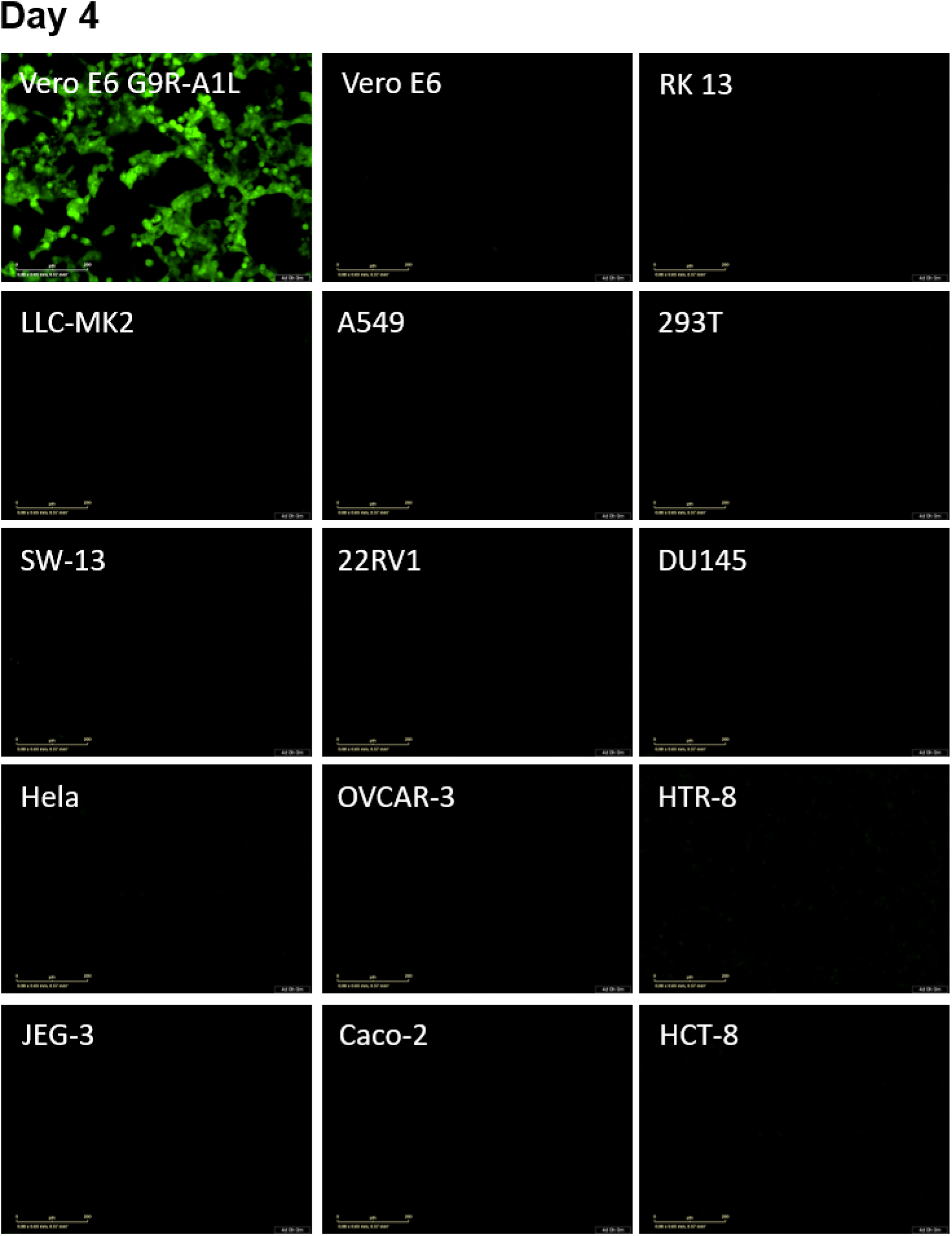
Analysis of deficient MPXV-infected non-complementing and complementing cell lines using fluorescence microscopy. 1 MOI of MPXV G9R+A1L-KO virus was inoculated onto 14 non-complementing cell lines. The Vero E6 G9R+A1L cell was served as a positive control. Representative mNeonGreen positive images at 4 days post infection are shown. Scale bar, 200 µm. Vero E6, African green monkey kidney cells; RK-13, rabbit kidney cells; LLC-MK2, rhesus kidney cells; A549, human non-small cell lung cancer cells; 293T, human embryonic kidney cells; SW-13, human adrenocortical small cell carcinoma cells; 22RV1, human prostate cancer cells; DU 145, human prostate cancer cells; Hela, human cervical cancer cells; OVCAR-3, human ovarian cancer cells; HTR-8, human chorionic trophoblast cells; JEG-3, human chorionic carcinoma cells; Caco-2, human colorectal adenocarcinoma cells; HCT-8, human ileocecal cancer cells.

**Extended Data Figure 4.**
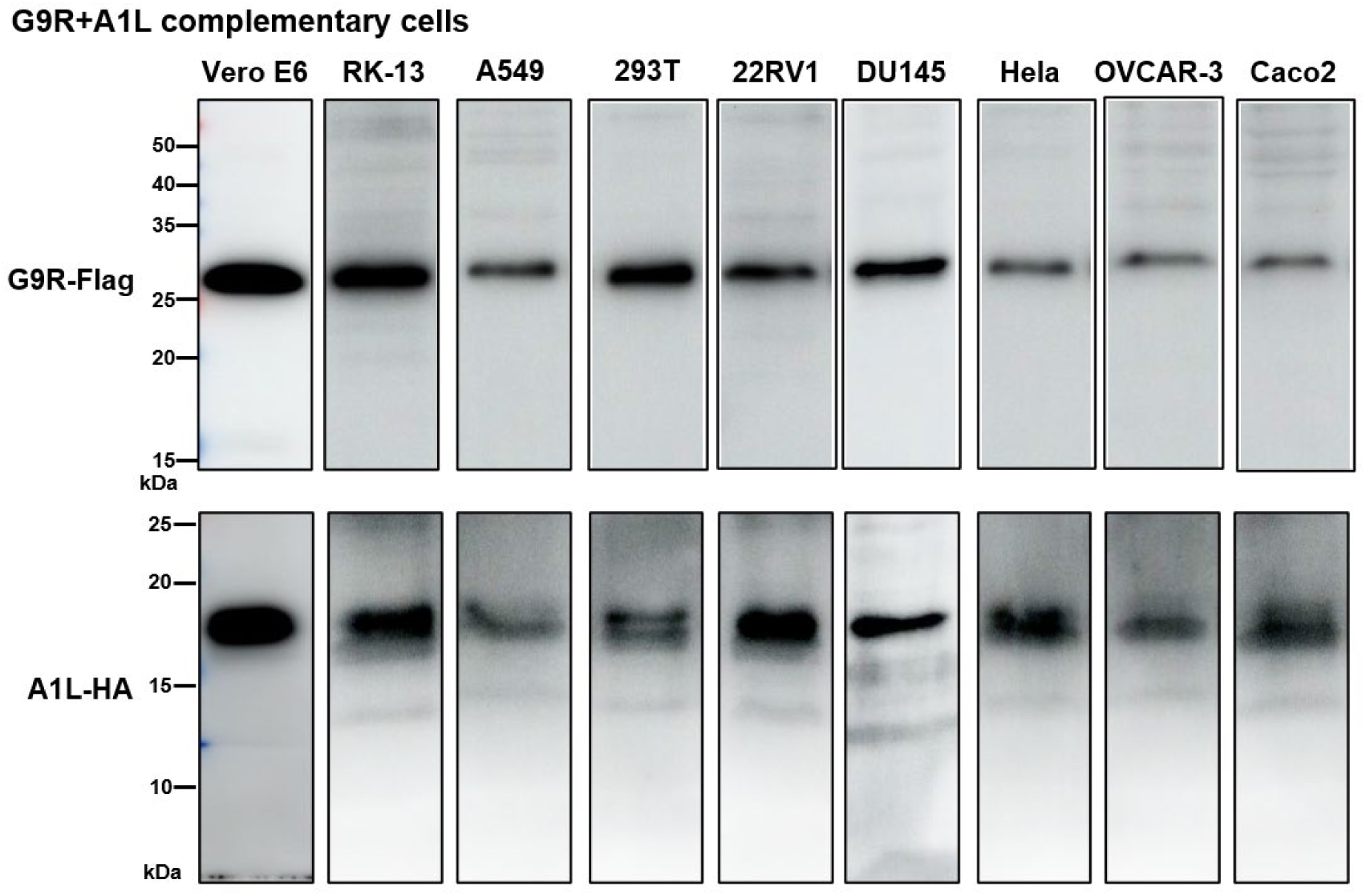
The expression of G9R and A1L genes in various complementary stable cell lines. The expression of G9R and A1L genes in 9 G9R+A1L complementing cells were detected by western blot. The G9R and A1L proteins were respectively fused with flag or HA tags at the N-terminal of the proteins. For different types of cells, the cell numbers for the blotting are normalized to an equal amount.

**Extended Data Figure 5.**
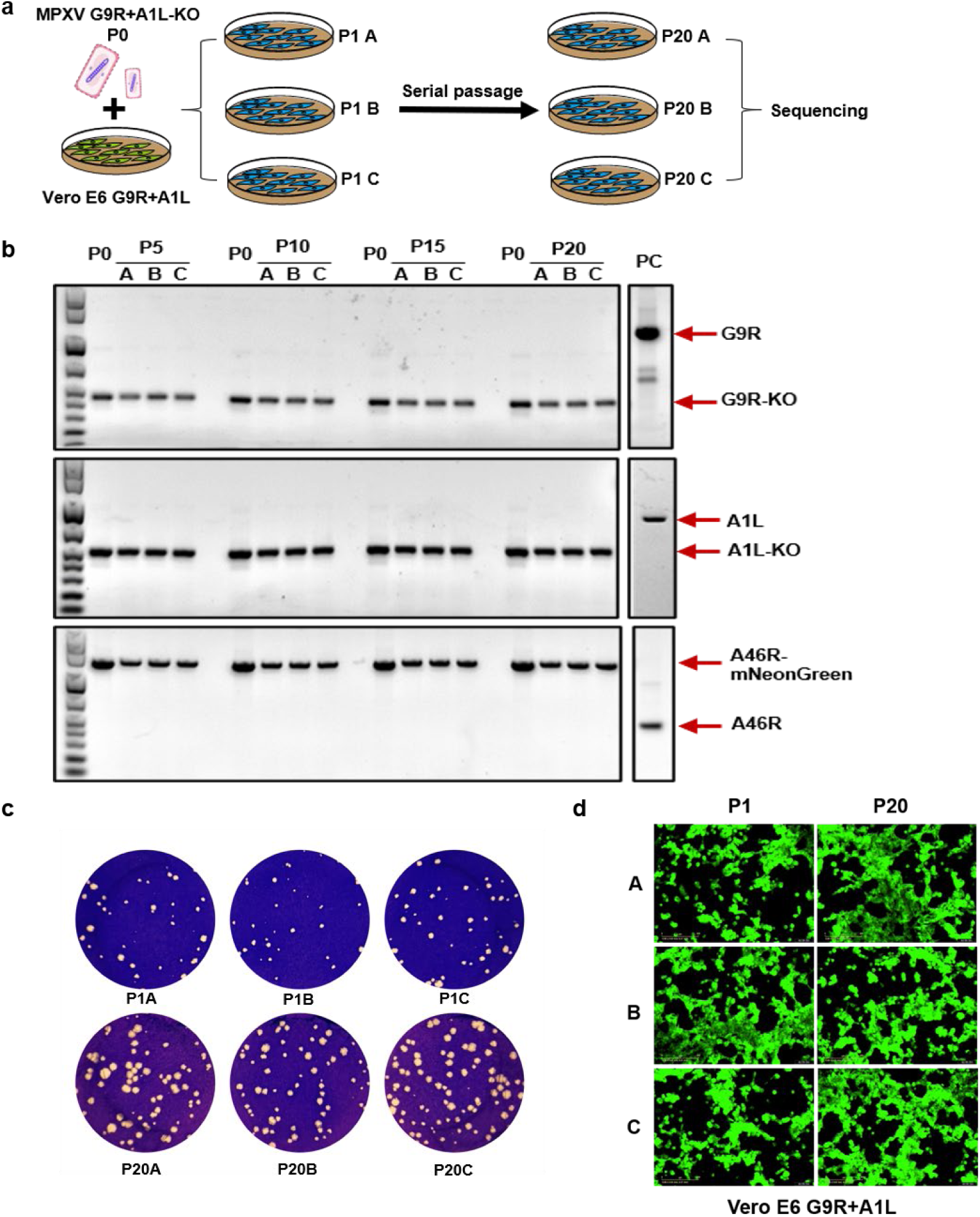
The serial passage of deficient MPXV in Vero E6 complementing cells. **a,** Experimental scheme of serial passage of deficient MPXV in Vero E6 G9R+A1L cells. MPXV G9R+A1L-KO virion was continuously passaged on Vero E6 G9R+A1L cells for 20 rounds in 3 replicates. The full-length MPXV genome of P0 and P20 were subjected to sequencing. **b,** PCR confirmation of G9R and A1L gene deletions and mNeonGreen insertion in MPXVs across various passages. The samples were collected at passages 5, 10, 15, and 20, with P0 initial viruses serving as controls. **c,d** The plaque morphology (**c**) of fluorescence (**d**) comparison of deficient MPXV after serial passages. **c,** The plaques of MPXV G9R+A1L-KO virus at passage 1 and passage 20 in Vero E6 G9R+A1L cells were stained with crystal violet 6 days post-infection. **d,** Vero E6 G9R+A1L cells were infected with P1 and P20 MPXV deficient viruses at MOI of 0.1. Representative mNeonGreen positive images are shown 4 days post-infection. Scale bar, 200 µm.

**Extended Data Table 1.** The detected mutations of deficient MPXV after 20 passages.

| No. | Genome region* | Product | Nucleotide | Amino acids | Replication |
| --- | --- | --- | --- | --- | --- |
| 1 | ITR | Noncoding Region | ^627T | N/A | A,C |
| 2 | MPXVgp016 | D13L | G18318A | silent mutation | A |
| 3 | MPXVgp076 | G7R | C71481T | S32L | C |
| 4 | MPXVgp087 | L3R | C79608T | silent mutation | A,C |
| 5 | Between MPXVgp125 and gp126 | A15.5L | ^121859T | Encode differently after K29 | A |
| 6 | MPXVgp149 | A39R | Δ143739C | frame shift | B |
| 7 | MPXVgp149 | A39R | T143761G | L79R | C |
| 8 | MPXVgp149 | A39R | G144060T | D179Y | A |
| 9 | MPXVgp150 | A40L | G145081T | Q113K | A |
| 10 | MPXVgp151 | A41L | C146786A | frame shift | A,C |
| 11 | Between MPXVgp151 and gp152 | Noncoding Region | Δ146878-146922 | N/A | A,B,C |
| 12 | MPXVgp152 | A42R | Δ146999T | frame shift | A,B,C |
| 13 | Between MPXVgp172 and gp173 | Noncoding Region | T170970G | N/A | A,B,C |
| 14 | Between MPXVgp179 and gp180 | Noncoding Region | Δ179151-179204 | N/A | A,B,C |
| 15 | ITR | Noncoding Region | ^196594T | N/A | A,C |

The mutations in the noncoding and coding regions of MPXV G9R+A1L-KO between passages P0 and P20 are noted. \* symbol means the region of MPXV MA001 genome (ON563414.3), ^ symbol means insertion, Δ symbol means deletion. A or B or C is biological replicate; N/A, Not Applicable. Sequences have been deposited to figshare website (Licence ID: CC BY 4.0, DOI: 10.6084/m9.figshare.26340355).

**Extended Data Table 2.** The IC_50_ values derived from plaque formation assay and mNeonGreen positive cell counting assay.

| Drugs | Cells | IC<sub>50</sub> uM (PFU) | IC<sub>50</sub> uM (Fluorescence) |
| --- | --- | --- | --- |
| Tecovirimat | Vero E6 G9R+A1L | 0.0038 | 0.002215 |
| Cidofovir | Vero E6 G9R+A1L | 6.776 | 1.239 |
| Raltitrexed | A549 G9R+A1L | 0.0823 | 0.2542 |
| Tecovirimat | A549 G9R+A1L | 0.0652 | 0.01138 |
| Cidofovir | A549 G9R+A1L | 48.29 | 12.83 |
| Brequinar | A549 G9R+A1L | 0.1236 | 0.4039 |
| Buparvaquone | A549 G9R+A1L | 3.854 | 5.956 |
| Narasin | A549 G9R+A1L | 0.0583 | 0.1294 |

**Extended Data Table 3.**
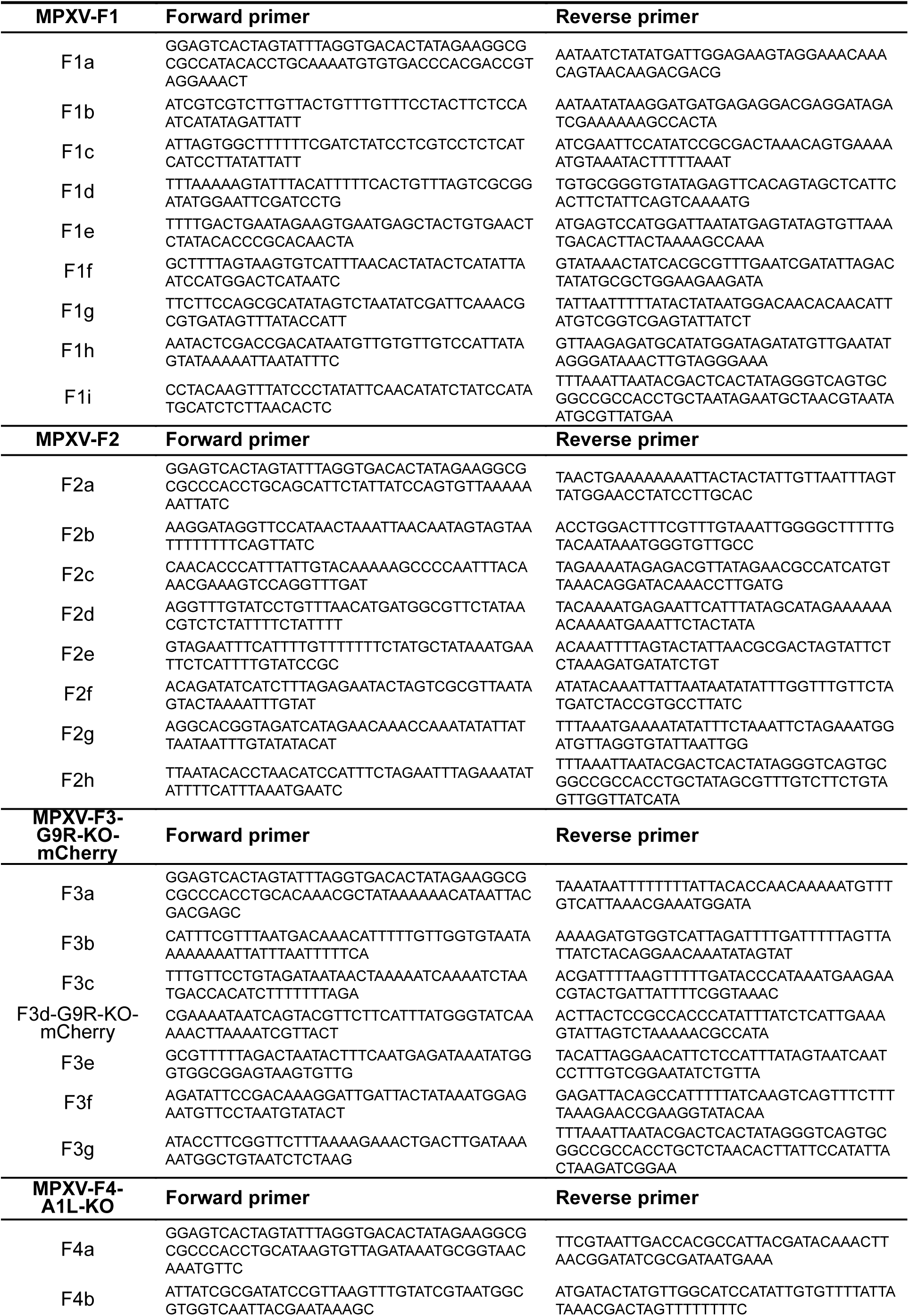
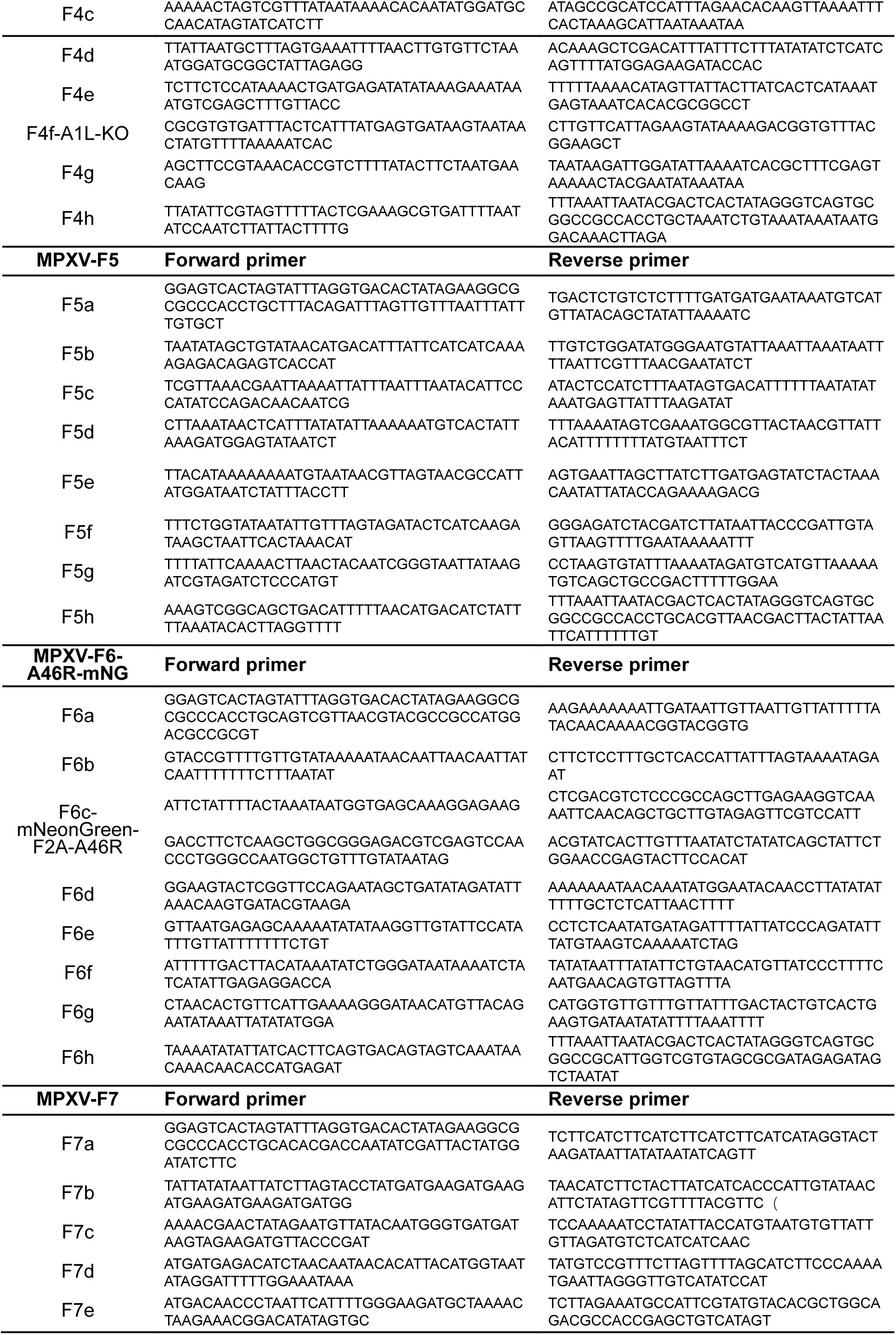
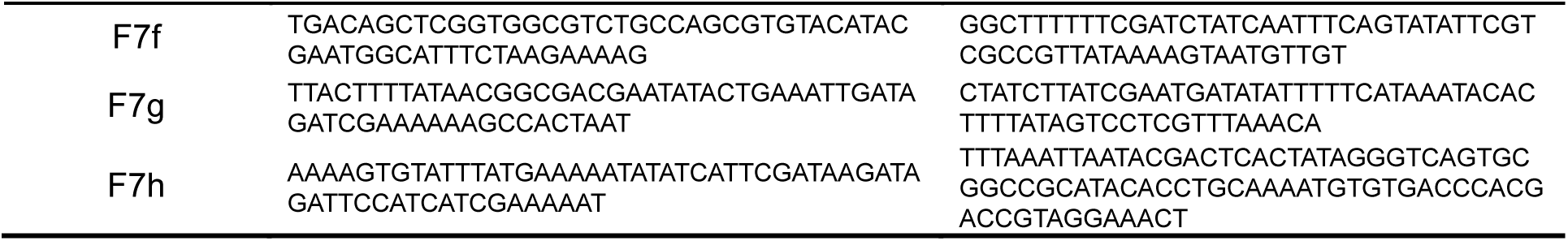
Primers used for construction of deficient MPXV.

